# Possible role of accessory proteins in the viral replication for the 20I/501Y.V1 (B.1.1.7) SARS-CoV-2 variant

**DOI:** 10.1101/2021.04.30.442222

**Authors:** Dimpal A. Nyayanit, Prasad Sarkale, Anita Shete-Aich, Abhinendra Kumar, Savita Patil, Triparna Majumdar, Shreekant Baradkar, Pranita Gawande, Sreelekshmy Mohandas, Pragya D Yadav

**Author notes:** **Corresponding author** Dr Pragya D. Yadav, Scientist ‘E’ and Group Leader, Maximum Containment Facility, Indian Council of Medical Research-National Institute of Virology, Sus Road, Pashan, Pune, Maharashtra, India Pin-411021.

## Abstract

The study investigates the replication cycle and transcriptional pattern of the B.1.1.7 variant. It was observed that the B.1.1.7 variant required a longer maturation time. The transcriptional response demonstrated higher expression of ORF6 and ORF8 compared to nucleocapsid transcript till the eclipse period which might influence higher viral replication. The number of infectious viruses titer is higher in the B.1.1.7, despite a lesser copy number than B.1, indicating higher infectivity.

## Introduction

The recent identification of the new Severe Acute Respiratory Syndrome Coronavirus-2 (SARS-CoV-2) variants has been of public health concern globally, after its first report from China in December 2019 (Zhu et al., 2020). SARS-CoV-2 encodes for structural [E (envelope), N (nucleocapsid), M (membrane), and S (spike)], a large polypeptide cleaved into different non-structural proteins and six accessory proteins (ORFs 3a, 6, 7a, 7b, 8, 10) (Zhou et al., 2020). The expression of these proteins is responsible for the replication of the SARS-CoV-2 with its hosts.

Information on the growth kinetics helps in understanding the virus-host interaction. It is also known that like other RNA viruses, The mutation rate of SARS-CoV-2 is higher in comparison to other virus groups (Duffy, 2018). The growth kinetic study of the SARS-CoV demonstrated the viral release at 7th-hour post-infection (hpi) in the Vero-E6 cells (Keyaerts et al., 2005). The growth kinetics study for the SARS-CoV-2 demonstrates a similar trend to SARS-CoV, where the virus was observed in the supernatant from 7^th^ hpi (Nyayanit et al., 2020). Further, limited data is present on the transcriptional response of the SARS-CoV-2 proteins after it infects a host cell (Blanco-Melo et al., 2020; Krishnamoorthy et al., 2021; Nyayanit et al., 2020; Sun et al., 2020) at different time-points. Comparison of respiratory viruses with the SARS-CoV-2 in cell lines and animal systems revealed a reduced cytokine and Interferon (Type I and III) response (Blanco-Melo et al., 2020). Transcriptome comparison of the SARS-CoV, MERS-CoV and the SARS-CoV-2 showed the suppression of the mitochondrial pathway and the glutathione metabolism (Krishnamoorthy et al., 2021). Time-point analysis by Sun et al demonstrated early host response to the SARS-CoV-2 entry to the host cell in comparison to SARS-CoV and MERS-CoV (Sun et al., 2020).

Delayed appearance of the cytopathic effect for the B.1.1.7 variant, with similar passage history, in comparison to the B.1 variant prompted us to investigate further. This study was carried out to understand the time required to complete the replication cycle of the B.1.1.7, a variant of concern (VOC). Further, in vitro transcriptional response generated at various time points was assessed.

### 20I/501Y.V1 (B 1.1.7) SARS-CoV-2 variant is a slow-growing virus

The intracellular live virus in pellet and extracellular live virus in supernatant was calculated in a triplicate set using endpoint titration assay in Vero CCL81 cells at defined time intervals. The presence of live virus was detected in pellet beginning from the 8^th^ hour’s post-infection (hpi) while the virus was released in the supernatant only at 12^th^ hpi (Figure 1). The time lag of 3h till the first virus appearance in the supernatant was observed. In the case of the B.1 variant, the virus was observed in the supernatant approximately within an hour after its appearance in the cell pellet (Nyayanit et al., 2020). The delayed observance of the B.1.1.7 in the supernatant in respect to B.1 led us to interpret that B.1.1.7 has a slow growth cycle caused due to its delayed maturation time.

**Figure 1.**
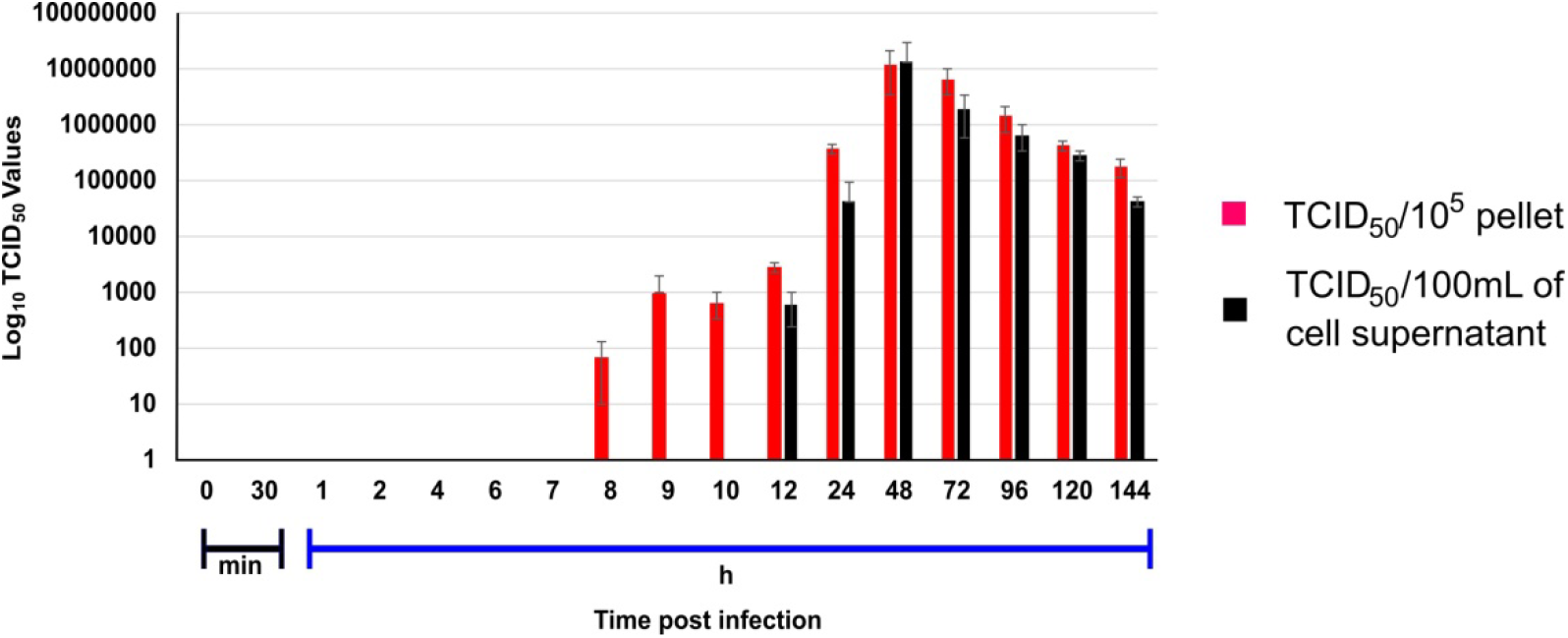
Growth kinetics of SARS-CoV-2 (20I/501Y.V1) in Vero CCL81 cell: Vero CCL81 cells were infected with 20I/501Y.V1 (0.1 MOI of 10^6^/ml TCID_50_) in triplicate sets. The cell pellet and supernatant were harvested at defined time intervals and viral titers were estimated using endpoint titration assay in Vero CCL81 cells. TCID_50_ of virus present within the cell pellet and in the supernatant is indicated with red and black colour respectively. Error bars depict the standard deviation observed in a triplicate set.

### Higher expression of accessory transcripts possibly leads to an increase in viral replication

*In vitro* transcriptional response for the SARS-CoV-2 genes was obtained using the next-generation sequencing of the cell pellet harvested at defined time intervals for the B.1.1.7 variant. The nucleocapsid transcript is expressed at a higher level in comparison to other viral transcripts at all-time points of experiments for SARS-CoV-2 (Finkel et al., 2020, Nyayanit et al., 2020). The RPKM values observed was hence normalized with the nucleocapsid expression. Figure 2A and B is the log10 plot of the normalized RPKM values obtained for structural protein transcript & non-structural polypeptide transcript and accessory protein transcript respectively at a defined time interval. The structural, non-structural, and accessory protein transcripts of the B.1 virus were always expressed in lower levels as compared to nucleocapsid transcript (Nyayanit et al., 2020). However, in the case of the B.1.1.7 variant, a few of the structural and accessory protein transcripts are expressed more than the nucleocapsid protein. The structural gene transcripts of B.1.1.7 are expressed in lower levels as compared to the nucleocapsid transcript at the time points (Figure 2A). But, the accessory protein ORF8 and ORF10 transcripts are expressed in higher levels at every time point in the study. The ORF6, ORF7a and 7b transcript are expressed at higher levels till the eclipse period of the virus and drops later. The expression of the ORF3a transcript is at lower levels for each time point (Figure 2B).

**Figure 2.**
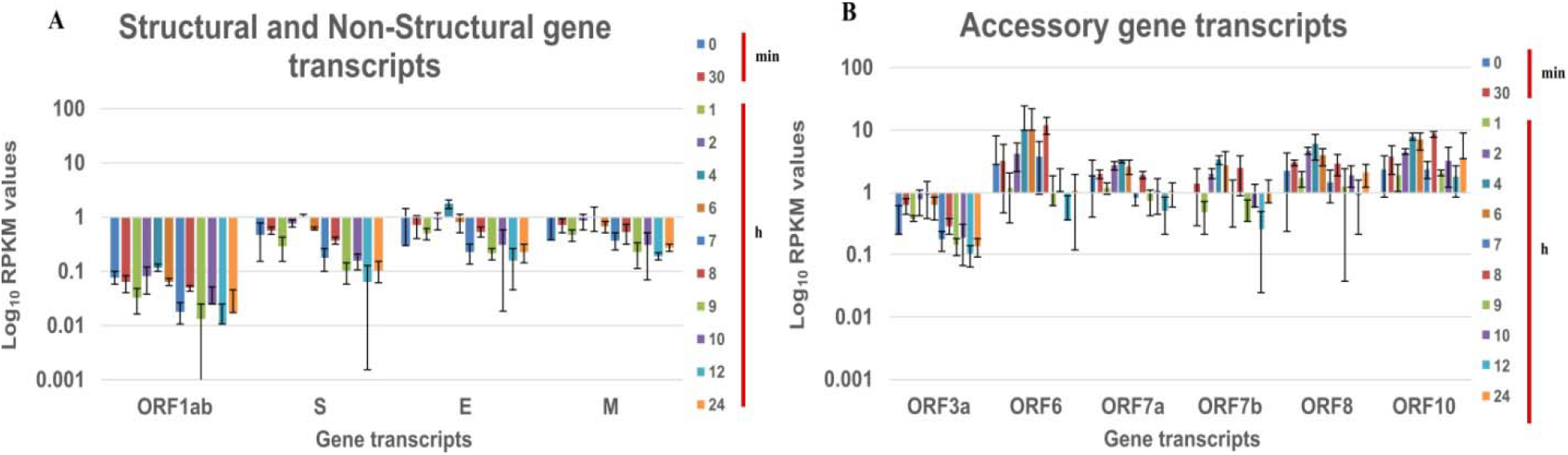
Expression of SARS-CoV-2 (20I/501Y.V1) genes in Vero CCL-81 at different time intervals in cell pellet: Normalized RPKM values of the SARS-CoV-2 (20I/501Y.V1) genes A) Structural and non-structural protein transcript B) Accessory protein transcripts from the cell pellet of Vero CCL-81 at different time intervals. The RPKM values are normalized with the nucleoprotein transcript values at the respective time intervals. Log10 RPKM values for the genes are plotted on the X-axis and gene transcripts on the Y-axis. Different colours are the log 10 RPKM values at that time intervals Average RPKM value is plotted and the standard deviation is shown in the error bar.

It has been reported earlier that pathogenic viruses increase viral replication by antagonizing the nuclear import to suppress the host immune response (Kato et al., 2021). ORF6 protein of the SARS-CoV-2 bind to the Nup98-Rae1 thereby targeting nuclear import (Miorin et al., 2020), indicated by its higher expression levels. Further, the role of ORF8 in viral replication was also reported (Muth et al., 2018). This indicates that the higher level of ORF6 and ORF8 can be associated with higher viral replication. With a similar observation in the increased expression level of the ORF6 and ORF8 transcript, it could be hypothesized that B.1.1.7 virus has higher viral replication.

### Higher transmission of B.1.1.7 variant can be possibly liked to larger infectious virus titer

The infectious virus titer of B.1.1.7 variant was ∼ 1×10^2.8^ per mL at 12 hpi whereas that of B.1 variant had ∼1×10^1.5^ per mL viral titer at 8 hpi when first detected in the supernatant. The sample for the B.1.1.7 was harvested on the 4^th^-day -post-inoculation (PID) whereas for B.1 it was the 3^rd^ PID. To understand the difference of infectious virus titre we infected both the strains to the Vero CCL81 cells and retrieved tissue culture fluids for the 3^rd^ and 4^th^ PID. It was observed that the infectious virus titer for B.1 variant was reduced by 30.8% on the 4^th^ PID. However, in the case of the B.1.1.7 variant, the infectious virus titre was reduced by 74.8% on the 4^th^ PID. This indicated that the amount of the infectious particle was less at the 4^th^ PID for B.1.1.7 variant in comparison to B.1 variant.

Further, the viral load of the B.1.1.7 variant was 7.86×10^11^ copies/ml at 4^th^ PID in comparison to the 3^rd^ 5.36×10^11^ copies/ml which is higher. An increase of ∼1.46 fold in the viral load was observed at the 4^th^ PID in comparison despite a 74.8% reduction in infectious virus particles reduced by titer. On the other hand, B.1 variant had ∼1.35 fold increase in viral load but, the infectious virus particles reduced only by 30.8% indicating a higher contribution of the live virus [3^rd^ PID: 1.69×10^12^ copies/ml and 4^th^ PID: 2.29×10^12^ copies/ml]. The real-time data indicates the B.1 variant to have a higher viral copy number. Overall higher viral copy number and a lesser reduction in viral titer were observed for the B.1 variant. The viral titer of the B.1.1.7 variant drops at a higher rate in the consecutive days however the amount of the infectious virus is still larger in comparison to the B.1 variant indicating replication. Further the ratio of the non-infectious: infectious is approximately in 3:1 ratio this indicates the infectious virus available for infecting a host cell is more in the B.1.1.7 variant. This makes it highly transmissible leading to a higher number of COVID-19 cases (Davies et al., 2021) but having reduced pathogenicity to a lesser infectious viral ratio.

This study reveals that B.1.1.7 SARS-CoV-2 variant is a slow-growing virus. Further, a higher replication along with reduced infectious viral titer was hypothesized to be linked to higher transmission with reduced pathogenicity.

## Materials and methods

The present study had approval by the Institutional Animal Ethics and Biosafety Committee of Indian Council of Medical Research (ICMR) -National Institute of Virology (NIV), Pune.

The 20I/501Y.V1 (B.1.1.7) SARS-CoV-2 isolate used in this study from the throat/nasal swab of United Kingdom traveller to India and submitted to GISAID (GISAID Number: EPI_ISL_825086) during late December 2020 (Yadav et al., 2021). Vero CCL81 grown in the 24-well plate was infected with the 20I/501Y.V1 SARS-Cov-2 strain (TCID_50_:10^5.5^/ml, Vero CCL81 P-3). A 0.1 multiplicity of infection (MOI) was used to study the viral replication and the transcriptional response of the host Vero CCL81 cell line in a triplicate set. Cell pellet and supernatant were harvested at the defined time intervals (0, 30 min, 1h, 2h, 4h, 6h, 7h, 8h, 9h, 10h, 12h, 24h, 48 h, 72h, 96h, 120h and 144h) and stored at -80°C till further use. TCID_50_ and Cyclic threshold (Ct) value for the *E* gene were determined calculated for both the supernatants and cell pellets using the end-point titration assay and real-time PCR method (Choudhary et al., 2020).

RNA was extracted from cell pellet and supernatant harvested at different time intervals. cDNA libraries were prepared for the extracted RNA and used for library preparation using the Illumina TruSeq Stranded mRNA LT Sample Preparation Kit. The libraries were sequenced using the Illumina platform. The total viral reads generated for the libraries of the harvested cell pellets and supernatants were mapped with the reference genome (accession number: NC_045512.1). The in vitro response for each gene transcript was quantified using the reads per kilobase million (RPKM) methods which normalizes the reads against the sequencing depth and gene length. The RPKM values were calculated using the CLC Genomics Workbench software (version 20.0.4, Qiagen Bioinformatics) for each time point. Further, the average of the observed RPKM values of each SARS-CoV-2 (20I/501Y.V1) genes transcripts was normalized against the nucleocapsid transcript at the respective time point.

## Acknowledgement

Authors gratefully acknowledge Prof. (Dr.) Priya Abraham, Director, ICMR-NIV, Pune. The author thanks the staff of Maximum Containment Facility, ICMR-NIV Pune especially Lt Col. (Dr.) Priyanka Pandit, Mr Rajen Lakra, Ms Manisha Dudhmal, Mrs Ashwini Waghmare, Kaumudi Kalele and Hitesh Dighe for technical support.

## Disclaimer

The findings and conclusions are of the authors, and the funding agencies have no role in any part of the study.

## Author Contributions

PDY contributed to study design, data analysis, interpretation and writing and critical review. DAN contributed to study design, data analysis and interpretation and writing. AMS, TP, SP, AK, PG and SM contributed to data collection and data interpretation. All the authors have read and approved the paper.

